# Germline *ERBB3* mutation in familial non-small cell lung carcinoma: expanding ErbB’s role in oncogenesis

**DOI:** 10.1101/2020.05.29.122796

**Authors:** Aideen M. McInerney-Leo, Hui Yi Chew, Po-Ling Inglis, Paul J. Leo, Shannon R. Joseph, Caroline L. Cooper, Satomi Okano, Tim Hassall, Lisa Anderson, Rayleen V. Bowman, Michael Gattas, Jessica E. Harris, Mhairi S. Marshall, Janet G. Shaw, Lawrie Wheeler, Ian A. Yang, Matthew A. Brown, Kwun M. Fong, Fiona Simpson, Emma L. Duncan

## Abstract

**Background:** Lung cancer is the commonest cause of cancer deaths worldwide. Although strongly associated with smoking, predisposition to lung cancer is also heritable with multiple common risk variants identified. Rarely, dominantly inherited non-small-cell lung cancer (NSCLC) has been reported due to somatic mutations in *EGFR/ErbB1* and *ERBB2.*

**Methods:** Germline exome sequencing was performed in a multi-generation family with autosomal dominant NSCLC, including an affected child. Tumour samples were also sequenced. Full-length wild-type (wtErbB3) and mutant ERBB3 (mutErbB3) constructs were transfected into HeLa cells. Protein expression, stability, and sub-cellular localisation were assessed; and cellular proliferation, pAkt/Akt, and pERK levels were determined.

**Results:** A novel germline variant in *ERBB3* (c.1946T>G: p.Iso649Arg), coding for receptor tyrosineprotein kinase erbB-3 (ErbB3), was identified, with appropriate segregation. There was no loss-of-heterozygosity in tumour samples. Both wtErbB3 and mutErbB3 were stably expressed. MutErbB3-transfected cells demonstrated an increased ratio of the 80kD form (which enhances proliferation) compared to the full-length (180kD) form. MutErbB3 and wtErbB3 had similar punctate cytoplasmic localisation pre- and post-EGF stimulation; however, EGFR levels decreased faster post-stimulation in mutErbB3-transfected cells, suggesting more rapid processing of the mutErbB3/EGFR heterodimer. Cellular proliferation was increased in mutErbB3-transfected cells compared to wtErbB3 transfection. MutErbB3-transfected cells also showed decreased pAkt/tAkt ratios and increased pERK/tERK 30 minutes post-stimulation compared to wtErbB3 transfection, demonstrating altered signalling pathway activation by mutErbB3. Cumulatively, these results support this mutation as tumorogenic.

**Conclusions:** This is the first reported family with a germline *ERBB3* mutation causing heritable NSCLC, furthering understanding of the ErbB family pathway in oncogenesis.

## INTRODUCTION

Lung cancer is the leading cause of cancer deaths worldwide (World Health Organisation) [1], with over 80% of cases attributable to smoking. However, lung cancer is also heritable, with heritability of ~18% [2]. Genome-wide association studies (GWAS) have identified multiple susceptibility loci for lung cancer overall (reviewed [3, 4]), for non-small cell lung cancer (NSCLC) [5] and for histology-specific sub-types of NSCLC [6] (with specific GWAS in squamous cell carcinoma [7] and adenocarcinoma [8], but not large cell to date). There have also been many reports of familial aggregation of lung cancer, (summarised [9]), with increased familial risk particularly observed in cases with younger age of onset [10, 11], of female gender, and with adenocarcinoma [12], even after adjusting for smoking status [12, 13]. Linkage and association studies in familial lung cancer have identified unique susceptibility loci, as well as confirming loci associated with NSCLC overall and with specific NSCLC subtypes [14–18]. Additionally, GWAS have identified unique susceptibility loci for NSCLC cases carrying somatic EGFR mutations [19, 20].

Somatic gain-of-function mutations affecting the tyrosine kinase (TK) domain of Epidermal Growth Factor Receptor (EGFR) are common in non-small cell lung cancer (NSCLC), particularly adenocarcinoma, and predict responsiveness to EGFR-targeting tyrosine kinase inhibitors (TKIs) [21]. Extremely rarely, germline carriage of *EGFR* mutations has been described in families with autosomal dominant NSCLC, occasionally with additional somatic *EGFR* mutations [22, 23]. EGFR (ErbB1, Human EGF Receptor [HER] 1) belongs to the ErbB family of receptor tyrosine kinases which includes ErbB2 (neu, HER2), ErbB3 (HER3) and ErbB4 (HER4). A germline *ERBB2* mutation was identified in another family with autosomal dominant NSCLC, without additional *HER2* somatic variant(s) [24]. No paediatric NSCLC were reported in these families; indeed, primary lung cancers in children are very rare [25, 26]. Notably, none of the loci associated with lung cancer in the many GWAS to date have included *EGFR* or other ERBB family members [4].

Here, we report a new causative gene in a family with autosomal dominant NSCLC.

## MATERIALS AND METHODS

This study was approved by The Prince Charles Hospital Metro North Human Research Ethics Committee (approval HREC/13/QPCH/216). Participants gave informed written consent.

Detailed methods are presented in Supplementary Data. Briefly, exome sequencing was performed on germline DNA in a multi-generational family with autosomal dominant NSCLC (Fig. 1). Given the rarity of autosomal dominant NSCLC, and paediatric lung malignancies overall [25, 26], analysis focussed on rare variants (previously unreported; and with minor allele frequency [MAF] <0.001), assessed against internal and external databases (e.g. gnomAD [27], 1000 Genomes [28], and dbSNP137 [29]).

**Figure 1.**
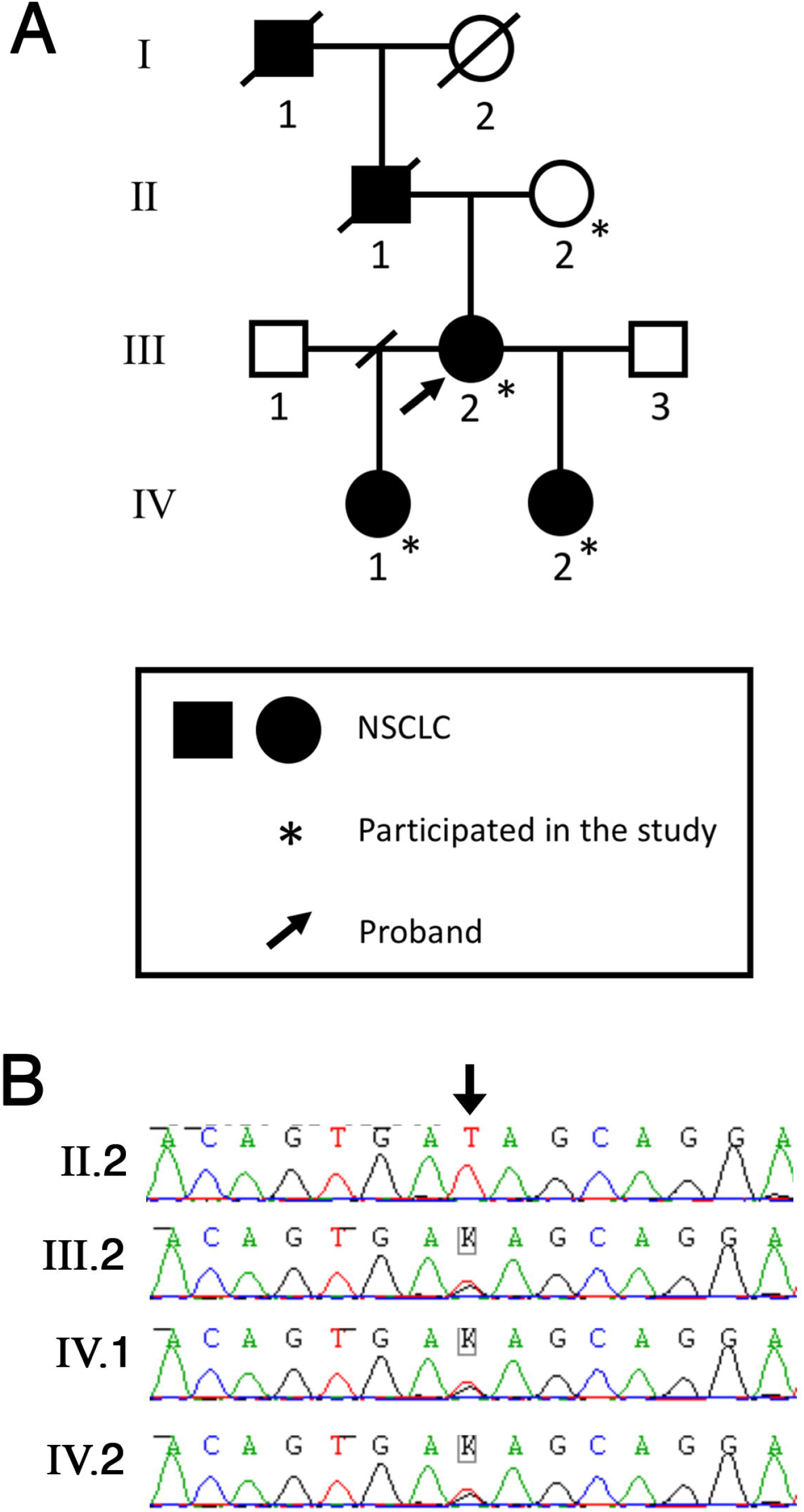
Germline *ERBB3* mutation segregating with NSCLC in an affected family. **A**. Family Pedigree. **B**. Sanger sequencing chromatograms of germline DNA, demonstrating heterozygosity for *ERBB3* c.1946T>G variant (arrow) in three affected individuals and wildtype in the proband’s unaffected mother.

Formalin-fixed paraffin-embedded [FFPE] samples were obtained from individuals LGCA-1.2, LGCA-1.3 and LGCA-1.6, with DNA extracted and sequenced. Expression and localisation of ErbB3 in normal and tumour tissue was assessed by immunohistochemistry.

Full-length wild-type (wtErbB3) and mutant (mutErbB3, c.1946T>G: p.Iso649Arg) *ERBB3* expression constructs were produced and transfected into HeLa cells (which do not express endogenous ErbB3 or ErbB2, but do express EGFR (ErbB1), the preferred dimerisation partner of ErbB3 [30]). To evaluate protein size and conformation, Western blotting was performed on lysates from transfected HeLa cells (vector-only, wtErbB3, or mutErbB3), probed with commercial anti-bodies against ErbB3 with β-tubulin used as protein-loading control. To assess localisation pre- and post-stimulation, transfected cells were either fixed (0’) or stimulated with 10ng EGF-Alexa Fluor 488 (30’) prior to fixation, and immunostained for ErbB3 and endogenous EGFR, with nuclei stained using DAPI. Transfected cells (vector-only, wtErbB3 or mutErbB3, co-transfected with green fluorescence protein) were separated by fluorescence-activated cell sorting and proliferation rate assessed. Signalling pathway activation of ErbB3 and mutErbB3 transfected cells were analysed by immunoblotting for ErbB3, EGFR, Akt (phospho- and total) and ERK (phospho- and total) in cells grown in full serum (control), 3 hours post-serum starvation (0) and post-EGF stimulation (10ng/ml) at 10 minutes and 30 minutes. Relative protein expression was quantified and the ratio normalised to β-Tubulin (used as a loading control) to enable the quantification of phospho-to total-Akt (pAkt/tAkt) and phospho-to total-ERK (pERK/tERK). Results are presented without formal statistical assessment, as is conventional for these analyses [31].

## RESULTS

### Clinical details

The proband (LGCA-1.2) presented with lung adenocarcinoma aged 51 years. Her father and paternal grandfather, died of NSCLC aged 39 and 34 years, respectively. Two of her five children have lung adenocarcinoma, presenting aged 12 and 30 years (Fig. 1). The proband, her father and grandfather had all smoked at some stage; however, neither of the children had ever smoked.

### Exome sequencing

Four novel good-quality variants affecting highly conserved bases and with appropriate familial segregation were identified, three of which were predicted damaging by at least two protein prediction algorithms (Table 1; Filtering steps presented in Supplementary Data: Table S1). Of these, the *ERBB3* variant (NM_001982; c.1946T>G; p.Ile649Arg) was of particular interest given the known oncogenic role of ErbB3 itself [32], and of other ErbB family members in heritable NSCLC In considering the other two variants: *SORBS1 (Sorbin and SH3 Domain Containing 1)* is involved in cell adhesion, growth factor signalling and cytoskeleton formation; but appears mainly to regulate insulin-mediated glucose uptake [33]. *ATG2B (Autophagy-Related Protein 2 Homolog B)* is involved in autophagy, a key pathway mediating stress-induced adaptation and cellular damage control. However, although exploited by cancer cells to survive stressors (e.g., starvation, hypoxia, and chemotherapy), autophagy is not considered an oncogenic driver process *per se* [34].

**Table 1.** Characteristics of Variants Fulfilling Filtering Criteria

| **Gene** | **Variant** | **GERP**<br>**score** | **SIFT** | **MutationTaster** | **PolyPhen2** |
| --- | --- | --- | --- | --- | --- |
| *ERBB3 (Erb-B2 Receptor Tyrosine Kinase 3)* | NM_001982:<br>c.1946T>G:<br>p.Ile649Arg | 3.87 | 0.00<br>(deleterious) | 0.995 (disease-causing) | 0.121 (benign) |
| *ATG2B (Autophagy related 2B)* | NM_018036:<br>c.4057T>A:<br>p.Cys1353Ser | 5.48 | 0.05<br>(deleterious) | 0.999 (disease-causing) | 0.978 (probably damaging) |
| *SORBS1 (Sorbin and SH3 domain-containing protein 1)* | NM_001034954:<br>c.2464A>G:<br>p.Ile822Val | 6.07 | 0.05<br>(deleterious*) | 0.999 (deleterious) | 0.903 (possibly damaging) |

Filtering the data with a less stringent MAF threshold (MAF<0.001) identified variants in eight additional candidate genes (Supplementary Data: Table S2), of which one *(PAXIP1)* is a genome stability gene previously associated with cancer [35]. Somatic copy number variation (CNV) of *PAXIP1* has also been associated with breast cancer prognosis [36]. However, the identified variant (rs199937188) is predicted benign and tolerated by Polyphen [37] and SIFT [38].

The data were also interrogated for coding variants in genes previously implicated in familial lung cancer (specifically, *EGFR, ERBB2, TP53* or *PARK2);* none were detected.

### Tumour sequencing

Sanger sequencing of tumour DNA excluded homozygosity of the *ERBB3* variant (data not shown). Unfortunately, tumour DNA from FFPE samples was too degraded for massively parallel sequencing, precluding comprehensive assessment of *ERBB3* somatic variants.

### Immunohistochemistry for ERBB3

ErbB3 is typically upregulated in NSCLC, staining both membrane and cytoplasm [39]. Tumour tissue from the proband (LGCA-1.2) showed weak cytoplasmic ErbB3, with absent staining of normal surrounding lung tissue (Supplementary Data: Figure S1) Results from other tumour samples were inconsistent; notably, less tissue was made available from these other individuals for this study, as their tumour samples were required to inform their ongoing clinical care.

### ErbB3 expression, folding, and cytoplasmic localisation

Both wtErbB3 and mutErbB3 were folded and expressed stably (Fig. 2A) with similar sub-cellular localisation (Fig. 2B). Cytoplasmic organelle distribution mirrored that of endogenous ErbB3 (in other, non-HeLa cells; data not shown). Compared with wtErbB3, cells expressing mutErbB3 showed a higher ratio of the 80kDa to full-length (~180kDa) forms (Fig. 2A). Without EGF stimulation, EGFR co-localised with mutErbB3 in concentrated puncta in the endosomal system, which was less evident with wtErbB3 (Fig. 2B). After EGF stimulation, both wtErbB3 and mutErbB3 increased in the perinuclear region (Fig. 2B), also co-localising with EGFR at this time-point (Fig. 2B, 30’ time-point).

**Figure 2.**
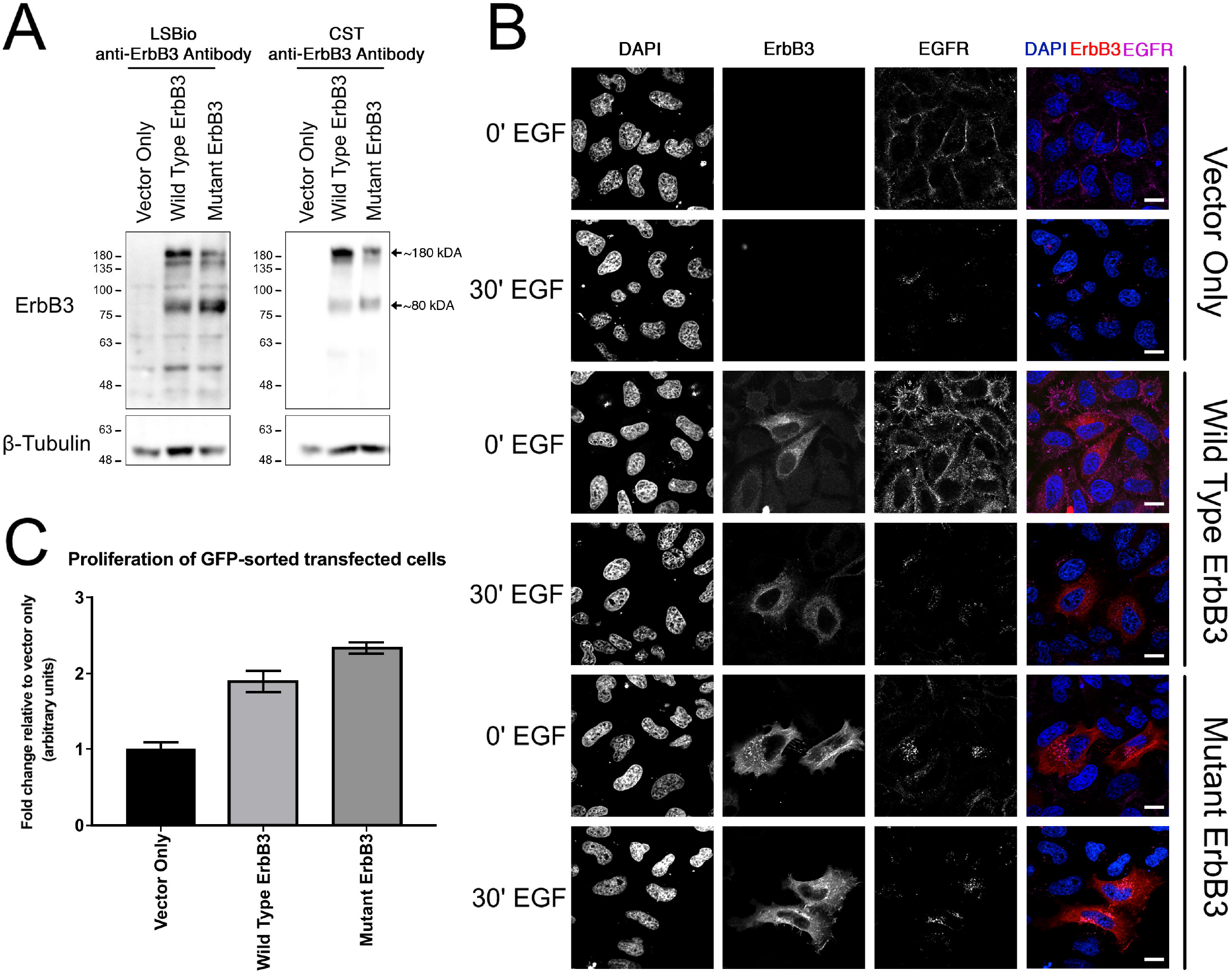
ErbB3 expression, folding, response to stimulation with EGF, and effect on cellular proliferation, in HeLa cells transfected with vector-only, wtErbB3, or mutErbB3. **A.** Western blot of lysates from transfected cells using two different commercial anti-ErbB3 antibodies. MutErbB3 is stably expressed and normally folded. A higher ratio of 80kDa to full length 180kDa form (arrows) is observed with mutErbB3 compared to wtErbB3. **B.** Transfected HeLa cells fixed pre- (0’) and post (30’)-EGF stimulation and immunostained for ErbB3 (red), endogenous EGFR (purple) and nuclei stained using DAPI (blue). Right column shows merged image. Scale bars, 20μm. Without stimulation EGFR co-localised with mutErbB3 in concentrated puncta, less evident with wtErbB3. After stimulation, both wtErbB3 and mutErbB3 increased in the perinuclear region, co-localising with EGFR. **C.** Proliferation assay of transfected cells quantified and described as fold change relative to vector only (data shown as mean ± S.E.M). MutErbB3-transfected cells showed increased rates of proliferation.

### Cell proliferation

HeLa cells expressing mutErbB3 demonstrated increased cellular proliferation, when compared to either HeLa cells expressing wtErbB3 or vector-only (Fig. 2C).

### Signalling Pathway Activation

After EGF stimulation, expression levels of mutErbB3 and wtErbB3 were comparable at all time-points; though, EGFR levels decreased over time in cells expressing mutErbB3 compared with wtErbB3 (Fig. 3A).

**Figure 3.**
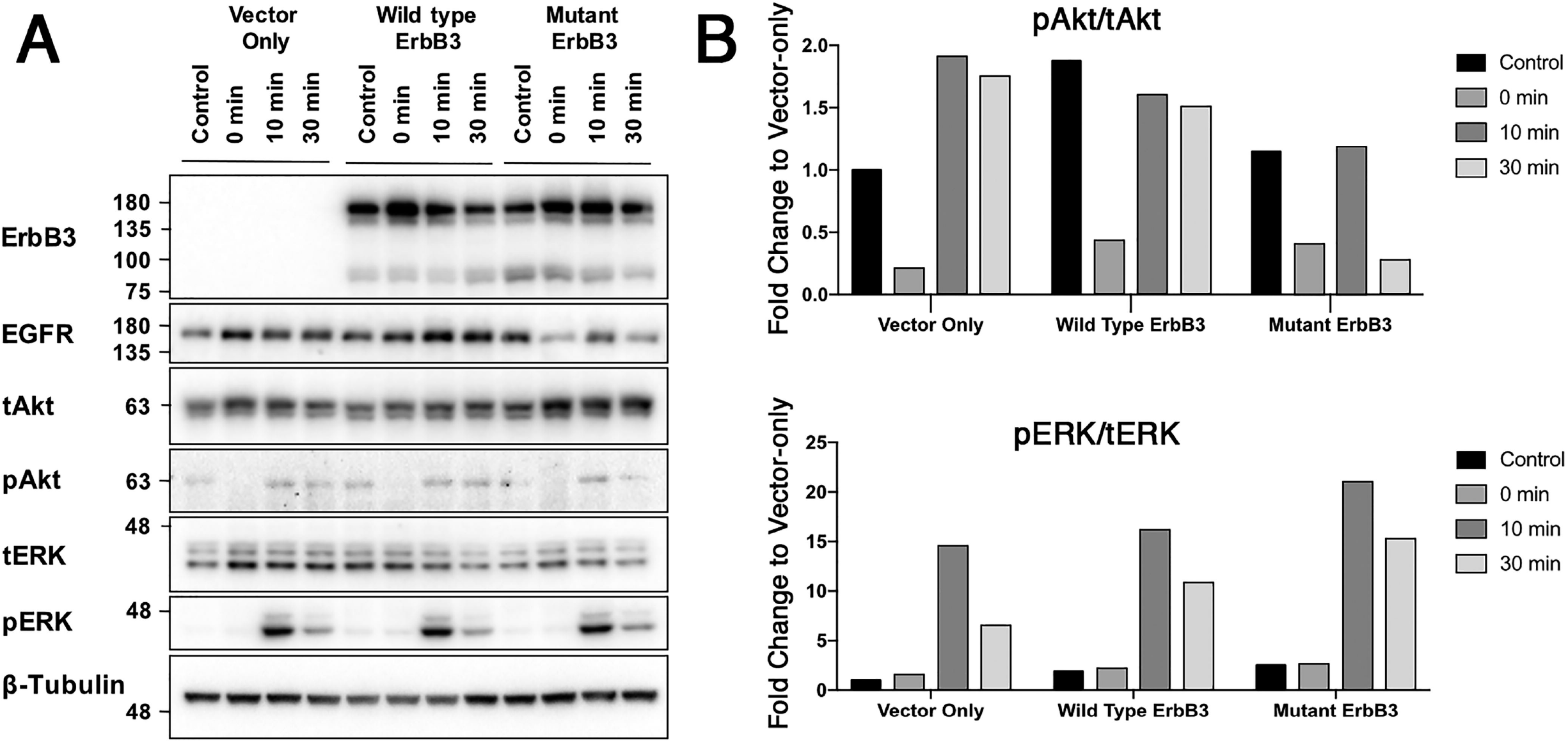
Analysis of protein expression and down-stream signalling pathway activation in HeLa cells transfected with vector-only, wtErbB3, or mutErbB3 constructs, before starvation (control), after 3 hours starvation (0 min) followed by EGF stimulation (assessed at 10 min and 30 min). **A.** Western blots of ErbB3, EGFR, phospho-Akt (pAkt), total Akt (tAkt), phospho-ERK (pERK) and total ERK (tERK), performed on transfected cell lysates. Following starvation and EGF stimulation, mutErbB3-transfected cells demonstrated decreased EGFR levels compared with wtErbB3. **B.** Ratios of phospho-Akt to total-Akt (pAkt/tAkt) (upper graphs), and phospho-ERK to total-ERK (pERK/tERK) (lower graphs) quantified and normalised to β-Tubulin. By 30 minutes, mutErbB3-transfected cells show decreased pAkt/tAkt ratio and increased pERK/tERK ratio compared with wtErbB3-transfected cells Both blot images and ratio quantification are representative of at least three separate biological replicates.

Cells expressing either mutErbB3 or wtErbB3 had increased pAkt levels, compared with vector-only transfected cells (Fig. 3B). After starvation followed by 10 minutes’ EGF stimulation, similar pAkt/tAkt ratios were observed in mutErbB3, wtErbB3, and vector-only transfected cells (Fig. 3B). However, by 30 minutes, mutErbB3-transfected cells had decreased pAkt/tAkt ratios compared with both wtErBB3 and vector-only transfected cells (Fig. 3B). Of note, mutErbB3-expressing cells had increased pERK at the 30-minute timepoint (Fig. 3B). These findings show that mutErbB3 is changing the signalling activation profile in response to ligand stimulation.

Together, these results suggest that the EGFR/mutErbB3 heterodimer is more efficiently activated, internalised and degraded, compared with EGFR/wtErbB3.

## DISCUSSION

We have identified a novel germline mutation in *ERBB3* segregating with autosomal dominant NSCLC. We demonstrate that mutErbB3 is stably expressed, functional with EGFR heterodimerisation and signalling, with an increased ratio of 80kDa vs. full-length 180kDa ErbB, a faster time-course of signalling activation and degradation, and increased cellular proliferation, compared to wtErbB3. These results support this mutation as the oncogenic driver of NSCLC in this family.

Multiple studies demonstrate the importance of ErbB3 in oncogenesis generally and NSCLC specifically [40]. *ERBB3* is part of a five-gene expression “signature” predictive of relapse-free and overall survival in NSCLC, independent of age, gender, stage and histological characteristics [41]. In a gene expression of ten “signature” genes in early lung adenocarcinoma, a two-gene signature comprising only *ERBB3* and *BRCA1* expression was an independent risk factor in predicting survival, improving the discriminatory power of conventional classification systems [42]. Other studies also identified increased ErbB3 expression correlating with shorter survival in NSCLC [43]. Within NSCLC, *ERBB3* expression is higher in adenocarcinoma compared with squamous [44] and other forms of lung cancer [45]; and circulating *ERBB3* mRNA levels correlate with higher TNM stage and poorer survival in adenocarcinoma [46].

The mutation reported here (c.1946T>G; p.Ile649Arg) lies in a conserved transmembrane motif, key to dimerization [47]. Notably the ERBB2 variant (p.Gly660Asp) previously associated with autosomal dominant NSCLC is located in the corresponding transmembrane motif of HER2 [24]. Although germline *ERBB3* variants have been reported previously [48], pathogenic variants have been reported extremely rarely – viz., a germline *ERBB3* mutation (c.4009G>A;p.Ala1337Thr), affecting the C-terminus of the protein, was reported in association with familial erythroleukemia [49]; and a homozygous loss-of-function mutation in *ERBB3* was associated with Lethal Congenital Contractural Syndrome Type 2 (OMIM 607598) in two Israeli-Bedouin families [50].

Although it has been hypothesised that germline polymorphisms in ErbB genes would contribute to lung cancer risk [51], no such associations have been identified in GWAS of lung cancer to date (neither lung cancer overall nor individual histopathological subtype) [4–8, 14]. Indeed, *ERBB3* has been ‘relatively under-investigated’ in lung cancer [51]. A very small single-candidate gene study suggested association of a variant in the *ERBB3* promoter region with lung cancer - but only with analysis restricted to a recessive model and limited to a non-smoking subset of 119 cases and 191 controls (*P*=0.037) [52]. Reduced ERBB3 expression was reported with the protective allele, consistent with an oncogenic role of ERBB3; however, these results have not been replicated in an independent cohort. Somatic *ERBB3* mutations, whilst common in colonic and gastric carcinomas, appear to be rare in NSCLC (Supplementary Data: Table S3). However, a study assessing CNVs in ErbB genes found that half of all lung adenocarcinomas have CNVs of *EGFR, ERBB2, ERBB3* and *ERBB4,* with higher CNV number corresponding to poorer prognosis [53].

Our germline *ERBB3* mutation is novel for NSCLC; and has not been reported (either as a somatic or germline mutation) in any other tumour type. Attributing causality to a variant segregating within a relatively small family just because of its rarity can lead to misattribution [54, 55]; hence our comprehensive functional assessment supporting this mutation as causative. Unsurprisingly, given the rarity of autosomal dominant NSCLC, no additional families were available for replication. However, our data concord with previous reports of germline mutations in ErbB family members *EGFR (ERBB1)* [56] and *ERBB2* [24] in other pedigrees with autosomal dominant NSCLC. Poor quality tumour DNA precluded assessment of *ERBB3* mutation(s) in our family, noting that somatic mutations were not identified in the single family with the germline *ERBB2* mutation and NSCLC [24], and inconsistently in individuals and families with *EGFR/ERBB1* mutations [56]. Although ErbB3 is expressed widely, this family has not manifested other malignancies (the proband has had non-cancerous colonic polyps). The apparent tissue specificity for malignancy is unclear, although again is consistent with NSCLC families with *EGFR* (ERBB1) [56] and *ERBB2* [24] mutations.

Our functional data support the identified variant as tumorogenic. Normally, ErbB3 (180kDa) is expressed as a transmembrane protein dimerised with another ErbB family member; upon activation, the heterodimer is internalised via ligand-induced receptor mediated endocytosis to endosomes and subsequently trafficked to lysosomes for proteolytical degradation. Additionally, some transmembrane ErB3 is directly cleaved, forming a cytoplasmic stable and active 80kDa form, which effects are normally offset by the tumour suppressor p14ARF sequestering the 80kDa form for degradation [57]. Our results suggest that mutErbB3 is more prone to cleavage, resulting in increased amounts of the cytoplasmic 80kDa form; moreover, this increase in the 80kDa form may exceed the sequestration capacity of p14ARF. Further, the 80kDa form may independently drive proliferation, as it can increase transcription of proliferative genes without requiring activation of cytoplasmic pathways [57]. Notably, our results demonstrated increased cellular proliferation in mutErbB3-transfected cells compared to wtErbB3.

We also demonstrated that mutErbB3 co-localised with EGFR; with EGFR levels decreasing over time in mutErbB3-transfected cells, compared to wtErbB3-transfected cells. Together, these results suggest that EGFR/mutErbB3 dimers internalise and reach the late endosomal/lysosomal system faster than wtErbB3-transfected cells, consistent with more rapid signal transduction with mutErbB3. Further, activation profiles of downstream signalling pathways differed in mutErbB3-transfected cells compared to wtErbB3-transfected cells; these pathways affect transcriptional regulation of cell proliferation and migration, both critical for cancer initiation and metastasis [32].

Our results may have clinical implications beyond genetic counselling. Both germline and somatic EGFR mutations affect NSCLC responsiveness to TKIs; and HER family-targeted therapy can induce prolonged progression-free survival specifically in individuals with TK domain mutations (including NSCLC) [58]. The identified *ERBB3* mutation does not lie within this domain; thus, HER family inhibitors may not benefit this family. However, ErbB3 downregulation (e.g. by siRNA) can restore tumour responsiveness to various therapeutic approaches, including TKIs, potentially of clinical relevance for this family [58].

**In conclusion,** we report the first family with heritable NSCLC segregating with a germline mutation in *ERBB3,* with functional data strongly supporting this mutation as oncogenic.

## Abbreviations

CNV: copy number variation;
EGFR: Epidermal Growth Factor Receptor;
GWAS: genome-wide association studies;
MAF: minor allele frequency;
NSCLC: non-smallcell lung cancer;
TKIs: tyrosine kinase inhibitors.

## Acknowledgements

We thank the family for their gracious participation. Additionally, we thank pathologist David Godbolt; research nurses Deborah Courtney, Linda Passmore, Elisabeth McCaul; Sharon Song for technical support; David Pennisi and Karolina Slater for writing and administrative support; and Malcolm Lim. The authors acknowledge the Translational Research Institute for providing the excellent research environment and microscopy and histology core facilities.

## Author Contributions

KF, RB, IY, JS, AML, ELD and MAB established the study. KF, PI, MG and TH identified and recruited family members to the study. ELD, AML, MB, JH, LA, PL, FS, SO, SRJ, CC and HYC designed and optimized experimental approach, performed the experiments and analyzed the data. ELD, AML and FS wrote the first draft of the manuscript, with additional input from, LW, HYC, SRJ and SO. All authors critically reviewed the final manuscript.

## Web resources

COSMIC: https://cancer.sanger.ac.uk/cosmic World Health Organisation statistics: http://globocan.iarc.fr/Pages/fact sheets cancer.aspx

**Supplementary Data**

