## supplemental data for "Germline *ERBB3* mutation in familial non-small cell lung carcinoma: expanding ErbB’s role in oncogenesis"

**SUPPLEMENTARY DATA**

**Supplementary Methods**

*Whole-exome sequencing*

DNA was extracted from blood or saliva using standard methods. DNA libraries were prepared using the TruSeq DNA v2 sample preparation kit (Illumina, San Diego, CA). Exome capture was performed using the Nimblegen SeqCap EZ Human v3.0 Exome Enrichment Kit (Roche, Basel, Switzerland). Library quality control and sample multiplexing were performed as previously described [1]. Massively parallel sequencing was performed using a HiSeq 4000, generating 100 base pair paired-end reads. Sample de-multiplexing, data alignment, duplicate removal, variant calling and annotation were as previously described [1]. After elimination of artefact and poor-quality variants, variants of potentially damaging consequence – "nonsynonymous single nucleotide variant (SNV)", "splicing", "frameshift", "stopgain SNV", "stoploss SNV" – were retained. Variant assessment used three protein prediction algorithms (SIFT [2], MutationTaster [3] and PolyPhen2 [4]). Variant minor allele frequency (MAF) was assessed against internal (>3000 exomes, performed with similar technologies) and external databases including gnomAD [5], ExAC [6], 1000 Genomes [7], and dbSNP137 [8]. Sequence data and results after filtering are shown in Supplementary Table 1

*Tumour sample DNA extraction and Sanger sequencing*

DNA was extracted from tumour samples by standard extraction kits. Sanger sequencing used the following primers:

F	CACTGGACACTTGGGACTCA

R	TGCACAGGGTACCACTCAAA

*Immunohistochemistry*

For immunostaining of ErbB3, FFPE tumour sections were subjected to antigen retrieval (EDTA 95°C 40m) in a decloaking chamber and stained with anti-ErbB-3/HER-3 Antibody (Clone 2F12; 1/300 O/N 4°C) for signal detection according to the manufacturer’s instructions (MACH 1 Universal HRP-Polymer Detection kit, Biocare Medical)). Negative controls included omission of the primary antibody and tissue negative for the protein.

*Antibodies for Immunohistochemistry*

Antibodies were purchased from commercial sources: mouse monoclonal anti-EGFR (31G7) (Life Technologies, San Francisco, CA); rabbit polyclonal anti-Akt, rabbit polyclonal anti-phospho-Akt (Ser473), rabbit polyclonal anti-P44/42 MAPK, and mouse monoclonal anti-phospho-P44/42 MAPK (Thr202/Tyr204)(p-ERK)(Cell Signalling Technology, Danvers, MA); mouse monoclonal anti-β-Tubulin (Invitrogen, Waltham, MA); rabbit monoclonal anti-HER3/ErbB3 (D22C5, CST, USA); mouse monoclonal anti-HER3/ErbB3 (LSBio,WA, USA) and mouse monoclonal anti-ErbB3/Her3 (Clone 2F12, Merck Millipore, Australia). All secondary antibodies were purchased from Invitrogen (Waltham, MA).

*Cloning protocol for ErbB3 mutant expression*

A clone was ordered from RIKEN (Wako, Japan), and sequence amplified by PCR. A construct designed to express the mutant ErbB3 (i.e., c.1946T>G) was engineered using Phusion Mutagenesis (Thermo Fisher Scientific, Waltham, MA), with the following primer sequences: Phos_T1964G_forward, GCTTTGACAGTGAGAGCAGGATTGGTAG (5´ phosphorylated) and Phos_T1964G_reverse, CATTGTCAGATGGGTTTTGCCGATCAG (5´ phosphorylated). Constructs with confirmed *ErbB3* mutant sequence were then amplified and purified for cell transfections. All procedures complied with physical and biological containment requirements in accordance with Office of the Gene Technology Regulator, Australian Government.

*Tissue culture*

HeLa cells were maintained in Dulbecco’s Modified Eagle’s Medium (Life Technologies, Thermo Fisher Scientific, Waltham, MA) supplemented with 10% heat-inactivated fetal bovine serum, 2mM of L-Glutamine and 10mM of HEPES. Cells were grown in a humidified incubator at 37°C with 5% CO_2_ level. Cell lines were SNV-confirmed and mycoplasma-tested monthly. Cells were transfected using Lipofectamine 2000 (Invitrogen, Waltham, MA). Transfection ratios were 2µg DNA/dish + Lipofectamine (1:2.5 ratio) for three hours in serum-free media prior to overnight incubation in full medium.

*Western blots*

Transfected cells were lysed with RIPA buffer (50mM Tris-HCl, pH 7.5/150mM NaCl/1% Triton X-100/0.1% sodium dodecyl sulphate (SDS)/0.5% sodium deoxycholate) supplemented with both protease and phosphatase inhibitors (Calbiochem, Merck KGaA, Darmstadt, Germany). Protein concentrations were determined by Pierce™ BCA Protein Assay (Thermo Fisher Scientific, Waltham, MA). 10μg of protein was separated on a 10% sodium dodecyl sulfate polyacrylamide gel electrophoretically and transferred to methanol-activated polyvinylidene fluoride membrane. Membranes were blocked with 2% bovine serum albumin (BSA) in TBS-T (20mM Tris-HCL, pH 7.6/137mM NaCl/0.1% Tween 20) for 1 hour. All primary antibodies were diluted in 2% BSA in TBS-T at 1:1,000. Membranes were incubated with primary antibodies overnight at 4°C with shaking before incubating with secondary antibodies at room temperature with shaking for 1 hour. Proteins were visualised using electrochemiluminescence (Thermo Fisher Scientific, Waltham, MA). Relative protein levels were determined and quantified using ImageJ software. Expression levels were normalised to β-Tubulin. All immunoblot images were processed identically using Adobe Photoshop software (Adobe Systems, San Jose, CA).

*Immunofluorescence*

HeLa cells were transfected with vector-only control, wild-type ErbB3 or mutant ErbB3 expression vectors. After 24 hours, cells were either fixed (0’) or stimulated with 10ng/mL of Alexa Fluor 488-conjugated EGF (Life Technologies, San Francisco, CA) (30’) prior to washing in PBS and fixation in 4% paraformaldehyde (PFA)/PBS for 20 minutes, followed by permeabilization in 0.1% TritonX100/PBS (PBTX) for 10 minutes and blocking in 2% (w/v) bovine serum albumin (BSA)/PBS. Coverslips were incubated for one hour in primary antibody (anti-EGFR), followed by washing in 2% BSA/PBS (Sigma-Aldrich)/PBS and incubation with secondary antibody (goat anti-mouse IgG Alexa Fluor^555^, Life technologies) and 30 minute DAPI staining. The coverslips were mounted on slides using ProLong® Gold antifade mountant (Life technologies). Images were acquired using a Zeiss 510 Meta confocal with a 63× objective.

*Epidermal Growth Factor (EGF) stimulation*

Cells were plated at 1×10^6^ cells/10cm dish and serum starved for three hours prior to EGFR being stimulated with 10ng/mL of Alexa Fluor 488-conjugated EGF (Life Technologies, San Francisco, CA). At 5, 15 and 30 minutes, cells were washed with ice-cold PBS before snap-freezing. Whole-cell protein lysates were obtained using RIPA buffer supplemented with protease and phosphatase inhibitor cocktail sets (Calbiochem, Merck KGaA, Darmstadt, Germany).

*Cell proliferation assays*

HeLa cells transfected with wild-type (wtErbB3) or Erb3 mutant (mutErbB3) expression vectors were harvested from a 10-cm dish before fluorescence-activated sorting for green fluorescent protein (GFP) cells. Sorted cells were washed once with media, and plated in a 96-well flat bottom plate at a density of 5,000 cells/well in 100μL media. The levels of metabolically active transfected cells were determined at two time-points: 24 hours and 48 hours after seeding. 20μL of CellTiter 96® AQueous One Solution Cell Proliferation Assay (Promega, Fitchburg, WI) was added to each well and the plate was incubated at 37°C for two hours, and absorbance measured at 490nm.

**Supplementary Results**

An average of 7.33Gb of sequence per individual was generated (mean depth of coverage, 54-fold).

**Table S1: Whole-Exome Sequencing Results Filtered for Novel Variants**

|  | III.2^#^ | IV.1^#^ | IV.2^#^ | II.2 |
| --- | --- | --- | --- | --- |
| Number of variants (MAF≤0.05) | 20,443 | 20,340 | 20,838 | 20,442 |
| Variants remaining after QC (including platform-related artefact) | 2,035 | 1,978 | 2,093 | 2,052 |
| Remaining variants of potentially damaging consequence* | 764 | 752 | 784 | 760 |
| Novel variants** | 84 | 92 | 115 | 96 |
| Variants affecting highly conserved bases regions*** | 43 | 50 | 68 | 59 |
| Heterozygous variant present in affected individuals only (i.e. absent in II.2) | 4 | | | |
| Variants predicted to be damaging by at least two protein prediction algorithms | Three variants identified (*ATG2B*, *ERBB3*, *SORBS1*) | | | |

MAF, minor allele frequency; QC, Quality Control

^#^ Lung adenocarcinoma cases

* Defined as nonsynonymous, frameshift insertions/deletions, stop/gain and splice site variants

** Not previously reported in internal or external databases, including ExAC, 1000 Genomes, and dbSNP137)

*** Genomic evolutionary rate profiling score (GERP) ≥2.5

**Table S2. Whole-Exome Sequencing Results Filtered for Variants with Minor Allele Frequency <0.001.**

|  | III.2^#^ | IV.1^#^ | IV.2^#^ | II.2 |
| --- | --- | --- | --- | --- |
| Number of variants (MAF≤0.05) | 20,443 | 20,340 | 20,838 | 20,442 |
| Variants remaining after QC (including platform-related artefact) | 2,035 | 1,978 | 2,093 | 2,052 |
| Remaining variants of potentially damaging consequence* | 764 | 752 | 784 | 760 |
| MAF <0.001** | 179 | 194 | 205 | 182 |
| Variants affecting highly conserved bases regions*** | 80 | 94 | 114 | 90 |
| Heterozygous variant present in affected individuals only (i.e. absent in II.2) | 12 variants identified (*CSRP3, MANSC4, RIMS4, DOCK7, NDUFAF5, MOB1B, ENPP6, PAXIP1, SRGAP1****, ATG2B****, SORBS1****, ERBB3****)* | | | |

MAF, minor allele frequency; QC, Quality Control

^#^ Lung adenocarcinoma cases

* Defined as nonsynonymous, frameshift insertions/deletions, stop/gain and splice site variants

** Not previously reported in internal or external databases, including ExAC, 1000 Genomes, and dbSNP137)

*** Genomic evolutionary rate profiling score (GERP) ≥2.5

****Identified as novel variants (see Table S1)

**Table S3: *ERBB3* somatic coding variants previously reported in NSCLC**

| **Cancer Cohort** | ***ERBB3* somatic variants in NSCLC** |
| --- | --- |
| TCGA | 19/1062 (11/531 adenoCa, 8/491 SCC) |
| COSMIC | 84/4398 (55/3172 adenoCa, 29/1226 SCC) |
| Jaiswal et al. [9] | 2/138 (1/71 adenoCa, 1/67 SCC) |
| Ding et al. [10] | 3/188 adenoCa (no SCC assessed) |
| Verlingue et al. [11] | 4/844 locally advanced, metastatic and/or irresectable cancers (adenoCa, SCC, or total NSCLC cases within this cohort unclear) |

TCGA, The Cancer Genome Atlas (https://portal.gdc.cancer.gov/genes/ENSG00000065361, accessed 15 October 2020); COSMIC, the Catalogue of Somatic Mutations in Cancer database, https://cancer.sanger.ac.uk/cosmic/search?query=ERBB3, accessed 15 October 2020). NSCLC, non-small cell lung cancer. adenoCa, adenocarcinoma. SCC, squamous cell carcinoma.

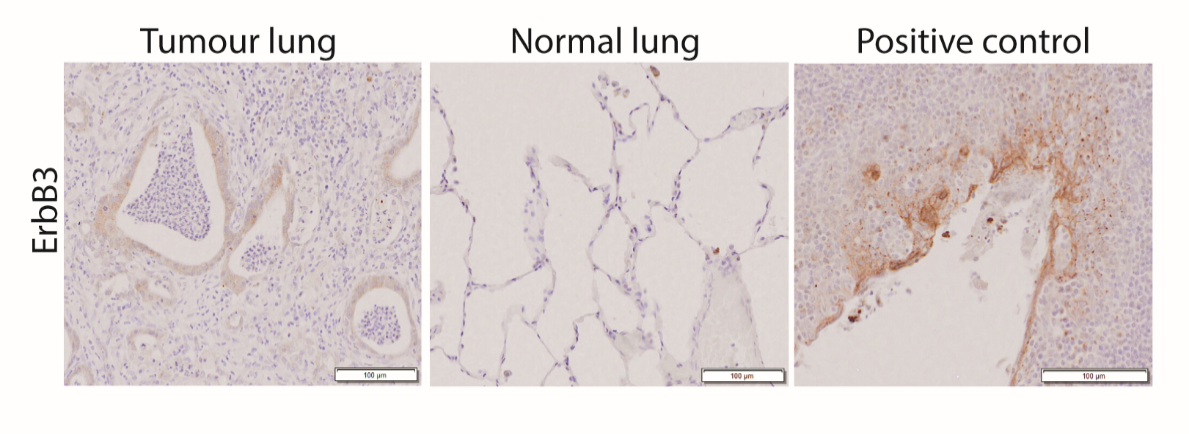

**Figure S1.** Immunohistochemical staining of ErbB3 in lung biopsy of proband (LGCA-1.2). Tumour tissue shows weakly positive (1+) cytoplasmic staining of ErbB3. Surrounding normal lung tissue was negative with macrophages showing moderately positive (2+) staining. Positive control tissue (tonsil) showing moderately positive (1-2+) membrane and cytoplasmic staining of ErbB3 in crypt epithelium. Scale bars: 100μm.
